# An evolutionarily conserved coreceptor gene is essential for CLAVATA signaling in *Marchantia polymorpha*

**DOI:** 10.1101/2021.02.22.427935

**Authors:** Go Takahashi, Shigeyuki Betsuyaku, Natsuki Okuzumi, Tomohiro Kiyosue, Yuki Hirakawa

**Author notes:** **Correspondence**: Yuki Hirakawa. These authors have contributed equally to this work (Go Takahashi, Shigeyuki Betsuyaku, Yuki Hirakawa).

## Abstract

Growth and development of land plants are controlled by CLAVATA3/EMBRYO SURROUNDING REGION-related (CLE) family of peptide hormones. In contrast to the genetic diversity of CLE family in flowering plants, the liverwort *Marchantia polymorpha* possesses a minimal set of CLE, MpCLE1(TDIF homolog) and MpCLE2 (CLV3 homolog). MpCLE1 and MpCLE2 peptides exert distinct function at the apical meristem of *M. polymorpha* gametophyte via specific receptors, MpTDIF RECEPTOR (MpTDR) and MpCLAVATA1 (MpCLV1), respectively, both belonging to the subclass XI of leucine-rich repeat receptor-like kinases (LRR-RLKs). Biochemical and genetic studies in Arabidopsis have shown that TDR/PXY family and CLV1/BAM family recognize the CLE peptide ligand in a heterodimeric complex with a member of subclass-II coreceptors. Here we show that three LRR-RLK genes of *M. polymorpha* are classified into subclass II, representing three distinct subgroups evolutionarily conserved in land plants. To address the involvement of subclass-II coreceptors in *M. polymorpha* CLE signaling, we performed molecular genetic analysis on one of them, Mp*CLAVATA3 INSENSITIVE RECEPTOR KINASE* (Mp*CIK*). Two knockout alleles for Mp*CIK* formed narrow apical meristems marked by _*prom*_Mp*YUC2:GUS* marker, which were not expanded by MpCLE2 peptide treatment, phenocopying Mp*clv1*. Loss of sensitivity to MpCLE2 peptide was also observed in gemma cup formation in both Mp*clv1* and Mp*cik*. Biochemical analysis using a *Nicotiana benthamiana* transient expression system revealed weak association between MpCIK and MpCLV1, as well as MpCIK and MpTDR. While MpCIK may also participate in MpCLE1 signaling, our data show that the conserved CLV3-CLV1-CIK module functions in *M. polymorpha*, controlling meristem activity for development and organ formation for asexual reproduction.

## Introduction

CLAVATA3/EMBRYO SURROUNDING REGION-related (CLE) peptides are a family of peptide hormones in land plants, mediating cell-to-cell communication in the plant body (Murphy et al., 2012; Hirakawa and Sawa, 2019; Fletcher, 2020). CLE peptides are genetically encoded as a precursor protein possessing a conserved CLE domain(s) at or near the C-terminus. Biosynthesis of CLE peptide hormone from the CLE domain involves post-transalational events including proteolytic cleavage, post-translation modifications and secretion to the apoplast (Ito et al., 2006; Kondo et al., 2006; Ohyama et al., 2009; Tamaki et al., 2013: Matsubayashi, 2014). In flowering plants, a large number of *CLE* genes are encoded in the genome, which have been extensively studied for the past two decades (Cock and McCormick 2001; Oelkers et al., 2008; Jun et al., 2010; Fletcher, 2020). The function of CLE genes cover a wide range of physiological processes including stem cell homeostasis in meristems, vascular cell differentiation, stomata differentiation and responses to various environmental cues (Fletcher et al., 1999; Suzaki et al., 2008; Okamoto et al., 2009; Stahl et al., 2009; Mortier et al., 2010; Etchels and Turner 2010; Hirakawa et al. 2010; Kondo et al., 2011; Fiume and Fletcher, 2012; Depuydt et al., 2013; Endo et al., 2013; Araya et al., 2014; Czyzewicz et al., 2015; Gutierrez-Alanis et al. 2017; Rodríguez-Leal et al., 2017; Qian et al., 2018; Takahashi et al., 2018; Ma et al., 2020). In bryophytes, which diverged early in the land plant lineage with respect to flowering plants (Puttick et al., 2018; Morris et al., 2018), relatively low number of *CLE* genes are encoded in the genome, providing simplified models to study the function of *CLE* genes (Bowman et al., 2017; Whitewoods et al. 2018). The minimal set of *CLE* genes, Mp*CLE1* (Mp6g07050) and Mp*CLE2* (Mp5g18050), are encoded in the genome of the liverwort *Marchantia polymorpha* (Bowman et al., 2017; Hirakawa et al., 2019; Montgomery et al., 2020; Figure 1A). MpCLE1 and MpCLE2 are the orthologs of TDIF (tracheary element differentiation inhibitor factor) and CLV3 (CLAVATA3) of *Arabidopsis thaliana*, respectively, representing the two distinct subgroups of CLE peptide family. In Arabidopsis, specific bioactivities of TDIF and CLV3 are attributed to the difference in a few amino acids between them, which are mediated by two distinct groups of receptors, TDIF RECEPTOR/PHLOEM INTERCALTED WITH XYLEM (TDR/PXY) and CLAVATA1/BARELY ANY MERISTEMs (CLV1/BAMs), respectively (Fletcher et al., 1999; DeYoung et al., 2006; Fisher and Turner, 2007; Ogawa et al., 2008; Hirakawa et al., 2008; Rodriguez-Villalon et al., 2014; Shimizu et al., 2015; Shinohara and Matsubayashi, 2015; Hirakawa et al., 2017; Crook et al., 2020). Since the ligand-receptor pairs are conserved among flowering plants and bryophytes and no CLE homologs were found in sister streptophyte algae, the specific CLE peptide-receptor pairs may have originated in the common ancestor of land plants (Whitewoods et al., 2018; Hirakawa et al., 2019; Hirakawa et al., 2020). In *M. polymorpha*, CLE genes regulate the activity of the apical meristem located at the apical notch of the thalloid gametophyte body. MpCLE1-MpTDR signaling acts as a negative regulator of cell proliferation at the apical notch, while MpCLE2-MpCLV1 signaling functions as a positive regulator of stem cell activity in the apical notch (Hirakawa et al., 2019; Hirakawa et al., 2020).

**Figure. 1.**
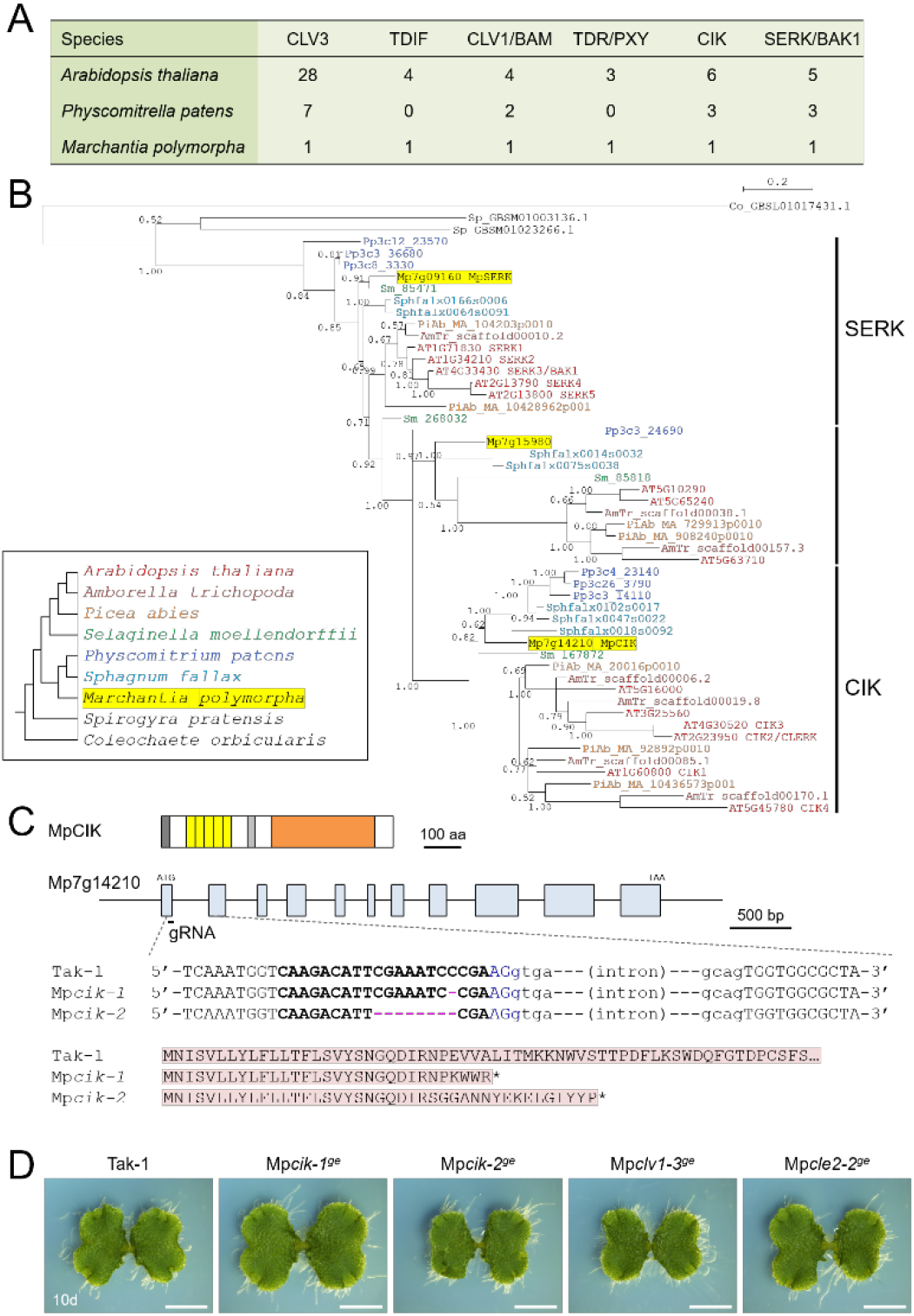
Analysis of LRR-RLK subclass II in *Marchantia polymorpha*. **(A)** The number of CLE-receptor homologs. **(B)** A phylogenetic tree of subclass-II LRR-RLKs, generated with a Bayesian method based on the conserved kinase domain. The posterior probabilities of trees are shown at the nodes. Coleochaete sequence was used as an outgroup. Land plant sequences form a monophyletic clade, which can be divided into three subgroups as indicated on the right. Inset shows the list of species with their phylogenetic relationships. **(C)** Gene/protein structures and genome editing alleles of MpCIK. (top) Protein structure of MpCIK. (middle) Structure of MpCIK/Mp7g14210 locus with the position of a designed guide RNA (gRNA). (bottom) Genotyping of genome editing alleles. Target guide sequence is in bold and PAM sequence is in blue. Deleted bases are indicated with hyphens in magenta. Exon and intron are indicated in capital and small letters, respectively. N-terminal region of WT and mutant proteins deduced from the genomic DNA sequences are indicated below. Asterisks indicate translational termination. **(D)** Overall morphology of 10-day-old plants grown from gemmae. Scale bars represent 0.5 mm.

Both TDR/PXY and CLV1/BAM belong to the subclass XI of leucine-rich repeat receptor-like kinase (LRR-RLK) family. In addition to CLE peptides, a number of peptide ligands have been shown to bind to specific members of subclass-XI receptors, which possess a long extracellular domain (ECD) composed of more than 20 LRRs (Shiu and Bleecker, 2001; Yamaguchi et al., 2006; Tabata et al., 2014; Ou et al., 2016; Shinohara et al., 2016; Song et al., 2016; Doblas et al., 2017; Nakayama et al., 2017; Doll et al., 2020). Accumulating evidence indicates that subclass-II receptors, such as SOMATIC EMBRYOGENESIS RECEPTOR KINASE/BRASSINOSTEROID INSENSITIVE1-ASSOCIATED KINASE1 (SERK/BAK1) family, participate in the peptide hormone perception by forming a heterodimeric complex with subclass-XI receptors (Hohmann et al., 2017; Gou and Li, 2020). Structural studies have revealed that SERK coreceptors have a short ECD containing five LRRs. The ECD of subclass-II receptors do not interact strongly or at all to the peptide ligand by themselves and rather recognize the ligand-receptor complex (Santiago et al., 2013; Sun et al., 2013; Wang et al., 2015; Santiago et al., 2016; Okuda et al., 2020). In line with this scheme, PXY/TDR and SERK2 are reported to form a heterodimeric complex for TDIF recognition, and multiple knockout mutants for Arabidopsis *SERK* genes show reduced TDIF sensitivity in vascular development (Morita et al., 2016; Zhang et al., 2016a; Zhang et al., 2016b).

In contrast to TDIF, involvement of SERK family has not been observed in CLV3-type CLEs. Instead, another group of subclass-II receptors, CLV3 INSENSITIVE RECEPTOR KINASEs (CIKs), have been implicated in CLV3 peptide perception. CIK proteins can form protein complexes with CLV1/BAM receptors (Hu et al., 2018; Cui et al., 2018). Quadruple mutants for Arabidopsis *CIK1-4* genes develop enlarged shoot apical meristems, which is similar to those of *clv* mutants. The growth from the enlarged meristems is not arrested by treatment with CLV3 peptide, a negative regulator of stem cells in Arabidopsis (Hu et al., 2018). Furthermore, full activity of Arabidopsis CLE26/CLE45 peptides in root phloem cell differentiation requires *CLE-RESISTANT RECEPTOR KINASE* (*CLERK*)/*CIK2* although biochemical interaction is not detected between the ECDs of CLERK and the subclass-XI receptor BAM3 (Anne et al., 2018). In this study, we searched for the homologs of *CIK* genes in *M. polymorpha* and analyzed their involvement in CLE peptide signaling by molecular genetic approach.

## Results

### A single CIK ortholog in *M. polymorpha*

In the *M. polymorpha* genome, three LRR-RLK genes (Mp7g09160/Mapoly0068s0069, Mp7g14210/Mapoly0009s0106, Mp7g15980/Mapoly0560s0001) have been classified into subclass II (Sasaki et al., 2007; Bowman et al., 2017; Montgomery et al., 2020). To better understand the evolutionary relationships, we performed phylogenetic analysis of the subclass-II genes from land plants (*Arabidopsis thaliana, Amborella trichopoda, Picea abies, Selaginella moellendorffii, Physcomitrium (Physcomitrella) patens, Sphagnum fallax, Marchantia polymorpha*) and charophycean algae (*Spirogyra pratensis, Coleochaete orbicularis*) based on the amino acid seuqnece of the kinase domain using a Bayesian method (Figure 1B). The tree inferred three subgroups diverged in the land plant lineage, each of which contains a single *M. polymorpha* gene. Mp7g14210, designated as Mp*CIK*, was grouped into a single subgroup with all *CIK* genes from Arabidopsis. Likewise, Mp7g09160/Mp*SERK* was grouped into the SERK subgroup with all Arabidopsis *SERK* genes. Thus, *M. polymorpha* genome may lack redundancy in CLE ligand/receptor/coreceptor orthlogs (Figure 1A).

### CRISPR-Cas9 editing of Mp*CIK* does not affect overall growth of gametophyte

To analyze the physiological function of the *CIK* coreceptor gene in *M. polymorpha*, we generated loss-of-function alleles for Mp*CIK* using CRISPR-Cas9 editing (Sugano et al., 2018). Sanger sequencing revealed that two independent transgenic lines, Mp*cik-1*^*ge*^ and Mp*cik-2*^*ge*^, possess different mutations at the CRISPR/Cas9 target site, both predetcted to result in gene knockout due to premature termination of translation (Figure 1C). We could not find significant differences in the overall morphology of thalli in 10-day-old Mp*cik-1*^*ge*^ and Mp*cik-2*^*ge*^ plants grown from gemmae, compared to any of wild-type (Tak-1), Mp*cle2-2*^*ge*^ and Mp*clv1-3*^*ge*^ genotypes (Figure 1D).

### Mp*CIK* is necessary for MpCLE2 signaling to control apical notch expansion

To analyze the involvement of Mp*CIK* in MpCLE2 peptide signaling, we examined the apical notch morphology in 4-day-old gemmalings grown on liquid M51C medium supplemented with or without 3 µM MpCLE2 peptide. In wild-type gemmalings, apical notches were expanded by treatment with MpCLE2 peptide, as reported previously (Figures 2A and 2B; Hirakawa et al., 2020). By contrast, apical notches in both Mp*cik-1* and Mp*cik-2* were insensitive to MpCLE2 peptide, which is similar to those in Mp*clv1-3* (Figures 2A and 2B). Importantly, apical notches of the Mp*cik* and Mp*clv1* alleles were narrower than those of wild type in the growth without MpCLE2 peptide, indicating that Mp*CIK* is involved in intrinsic MpCLE2-MpCLV1 signaling. Consistently, *Mpcle2-2*^*ge*^developed narrow apical notches but it was sensitive to the treatment with the MpCLE2 peptide as reported previously (Figure 2B; Hirakawa et al., 2020). _*pro*_Mp*YUC2(YUCCA2):GUS* is a marker for the tip of apical notch, and _*pro*_Mp*YUC2:GUS*-positive (Mp*YUC2*^*+*^) region is affected by MpCLE2-MpCLV1 signaling (Eklund et al., 2015; Hirakawa et al., 2020). Compared to wild-type (Tak-1) background, Mp*YUC2*^*+*^ region was reduced in Mp*cik* backgrounds, phenocopying Mp*clv1-3* (Figure 2C). These data support that MpCIK is an essential component of the MpCLE2 peptide perception.

**Figure 2.**
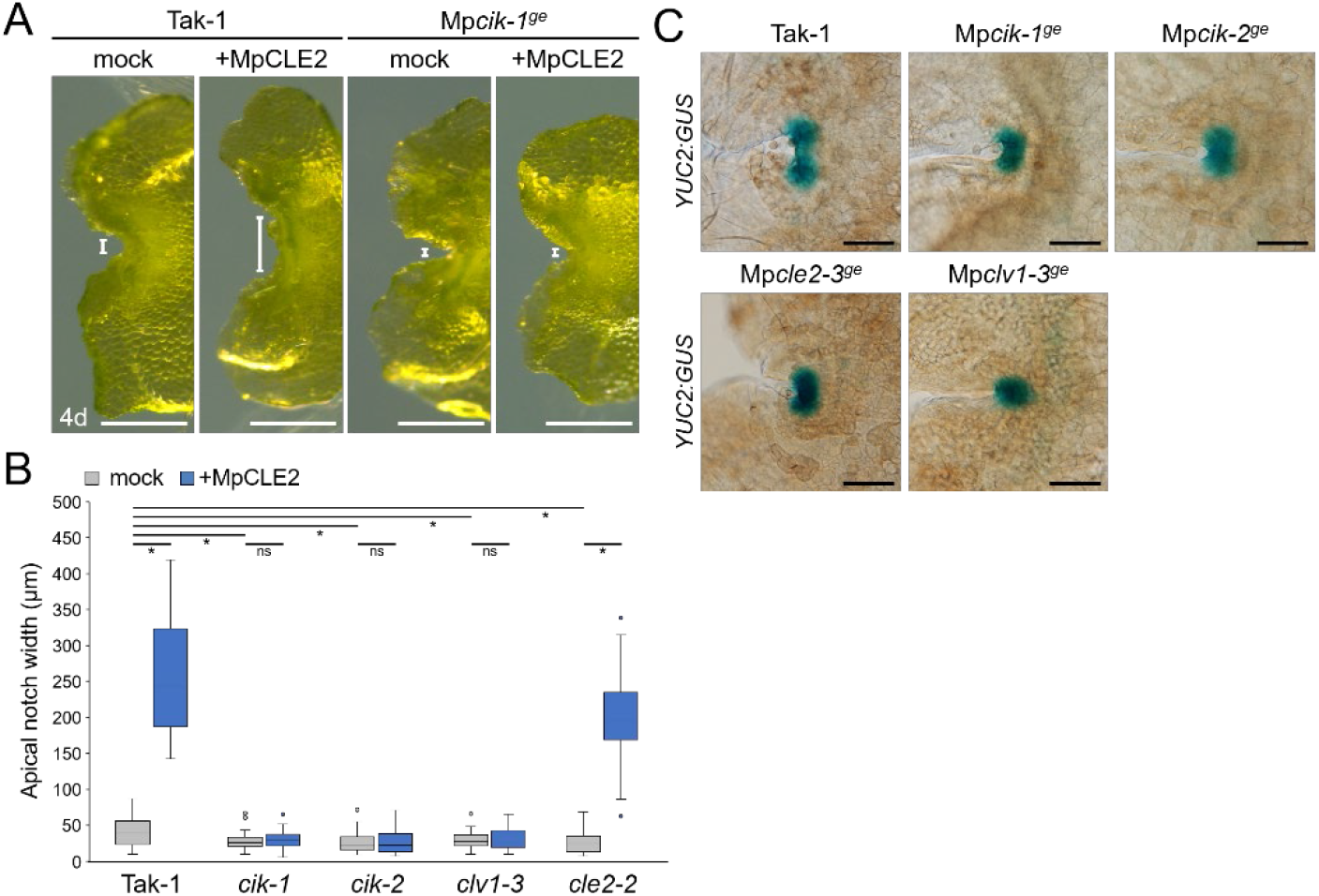
Phenotypes of Mp*CIK* knockout alleles in the apical notch. **(A)** Morphology of 4-day-old gemmalings grown with or without MpCLE2 peptide as indicated above. Width of apical notch is indicared by white lines. **(B)** Quantification of apical notch width. Asterisks indicate a significant difference in a two-tailed Welch’s *t* test, p < 0.05; ns, not significant; n = 30. **(C)** _*pro*_Mp*YUC2:GUS* marker in 4-day-old gemmalings. Genetic background is indicated above each panel. Scale bars represent 500 µm in (A) and 100 µm in (C).

### MpCLE2 signaling affects thallus branching and gemma cup formation via Mp*CLV1* and Mp*CIK*

To further address the involvement of Mp*CIK* in MpCLE2 signaling, we used _*pro*_Mp*YUC2:*Mp*CLE2*, a gain of function allele of Mp*CLE2*, which stably develops supernumerary branching from the expanded apical meristems (Hirakawa et al., 2020). We generated Mp*cik* and Mp*clv1* knockout alleles in _*pro*_Mp*YUC2:*Mp*CLE2-2* background by CRISPR/Cas9-mediated genome editing (Figure S1). Both Mp*cik-3*^*ge*^ and Mp*clv1-4*^*ge*^ suppressed the supernumerary branching phenotype in 16-day-old plants (Figure 3A), which is consistent with the results in peptide treatment assay (Figure 2B). Furthermore, the number of gemma cup formed on thalli was reduced in _*pro*_Mp*YUC2:*Mp*CLE2* and it was suppressed in both Mp*cik-3*^*ge*^ and Mp*clv1-4*^*ge*^ (Figure 3A). Time-couse analysis showed that the gemma cup formation was significantly reduced and delayed in _*pro*_Mp*YUC2:*Mp*CLE2* plants compared to wild type (Figure 3B). Meanwhile, all Mp*cik* and Mp*clv1* alleles showed minor increase in gemma cup formation compared to wild type (Figure 3B).

**Figure. 3.**
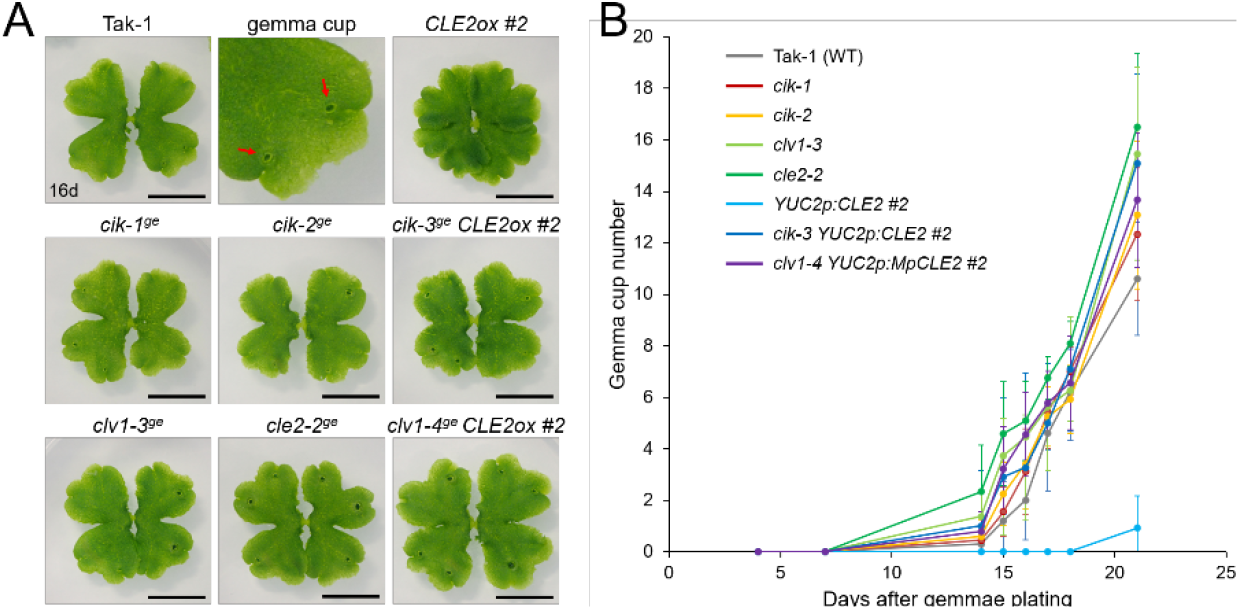
Mp*CIK* knockout suppresses gain-of-function phenotypes of Mp*CLE2*. **(A)** Overall morphology of 16-day-old plants grown from gemmae. Genotypes are indicated above the panels. The upper middle panel shows the magnification of Tak-1 image in which arrows indicate gemma cups. Note that _*pro*_Mp*YUC2:*Mp*CLE2* (*CLE2ox*) exhibits multichotomy and produces no gemmae cups at this age. Scale bars represent 1 cm. **(B)** Number of gemmae cups (mean and SD; n= 9-12). *CLE2ox* showed a significant delay of gemmae cup formation. Data are obtained at 4, 7, 14, 15, 16, 17, 18, 21 days after gemmae plating. This experiment was repeated twice with similar results.

### Biochemical interaction of MpCIK and MpCLV1 proteins

Since the genetic analysis suggests that MpCIK may function as a coreceptor for MpCLV1 for the perception of MpCLE2 peptide, we examined the biochemical interaction between MpCIK and MpCLV1 proteins expressed in a *Nicotiana benthamiana* transient expression system, which has been utilized to analyze the interaction of CLV and CIK receptors of Arabidopsis (Kinoshita et al., 2010; Betsuyaku et al. 2011; Hu et al. 2018). MpCIK, MpCLV1 and MpTDR were expressed under the control of 35S promoter in *N. benthamiana* as proteins C-terminally fused to 3× HAs-single StrepII or 3× FLAG (MpCIK-3HS, MpCLV1-3FLAG, MpTDR-3FLAG), respectively (Figure 4A). MpTDR, a receptor for MpCLE1, was also included in this interaction assay (Hirakawa et al. 2019). In co-immunoprecipitation experiments using an anti-HA affinity matrix, MpCLV1-3FLAG was detected not strongly but reproducibly in the immunoprecipitates containing MpCIK-3HS, suggesting a weak or transient interaction between MpCIK-3HS and MpCLV1-3FLAG (Figure 4B). Similarly, MpTDR-3FLAG was also shown to associate weakly with MpCIK-3HS (Figure 4B). Thus, MpCLV1 and MpTDR are capable of interacting with MpCIK.

**Figure 4.**
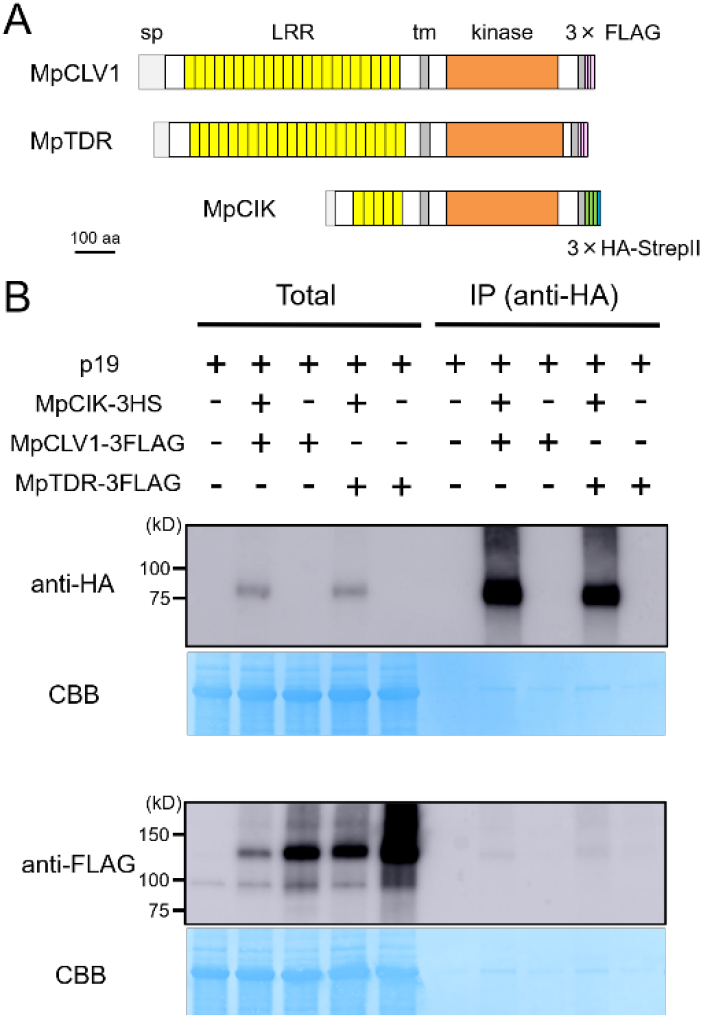
MpCIK weakly associates with MpCLV1 and MpTDR in *N. benthamiana*. (A) MSchematic illustration of expressed receptors. (B) Co-immunoprecipitation experiment using anti-HA affinity matrix. The indicated combinations of MpCIK-3HS, MpCLV1-3FLAG and MpTDR-3FLAG constructs, together with p19 silencing suppressor, were transiently expressed in *N. benthamiana*. Total proteins were extracted and immunoprecipitated with anti-HA affinity matrix. Immunoblot analyses were performed using anti-HA or anti-FLAG antibody. In the presence of MpCIK-3HS, MpCLV1-3FLAG and MpTDR-3FLAG were co-precipitated with anti-HA affinity matrix. This experiment was repeated twice with similar results.

### Mp*cik* is sensitive to MpCLE1 and TDIF

Our biochemical data suggests that MpCIK could also function as a coreceptor for MpTDR upon the perception of MpCLE1, a TDIF-type CLE peptide. Previously, we showed that synthetic TDIF-type peptides cause slight reduction of overall growth and twisted lobes in *M. polymorpha* thalli (Hirakawa et al., 2019). In order to address the possible involvement of MpCIK in MpCLE1 signaling, we first examined the effects of TDIF, the strongest analog among known TDIF-type CLE peptides including MpCLE1 peptide. In 14-day-old plants grown from gemmae on growth medium supplemented with 3 µM TDIF, both Tak-1 and Mp*cik-1* showed slight reduction of overall growth and twist in the thallus lobes (Figure 5A), indicating that Mp*CIK* is not necessary for the perception of TDIF. We further analyzed the effects of MpCLE1 overexpression using _*pro*_Mp*YUC2:*Mp*CLE1* transformants. In the wild-type (Tak-1) background, _*pro*_Mp*YUC2:*Mp*CLE1* resulted in small, twisted thalli in 14-day-old plants. This phenotype was not observed in _*pro*_Mp*YUC2:*Mp*CLE1*^*1-417*^, a truncated version of MpCLE1 lacking an essential asparagine residue in the CLE peptide motif (Figure 5B; Hirakawa et al., 2019). In the Mp*cik-1*^*ge*^ background, _*pro*_Mp*YUC2:*Mp*CLE1* also resulted in small twisted thalli although the effects on growth was mild compared to the Tak-1 background, as judged from the ground cover area in 14-day-old plants (Figures 5B and 5C). These data suggests that MpCIK could be partially involved in MpCLE1 perception.

**Figure 5.**
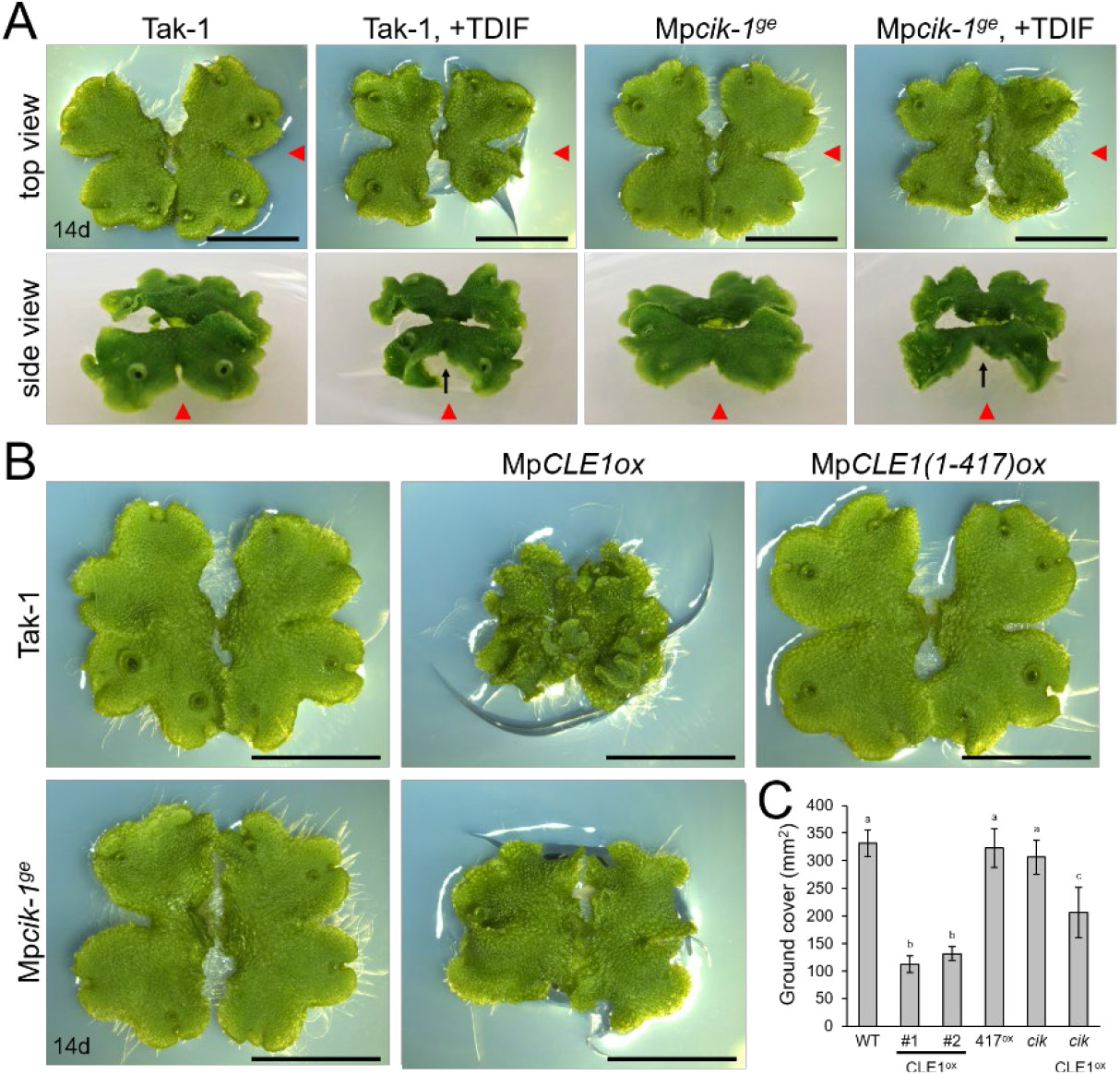
Growth of Mp*cik* thalli is sensitive to MpCLE1/TDIF activity. (A) Overall morphology of 14-day-old plants grown from gemmae on medium supplemented with or without TDIF peptide as indicated above. Side view panels show the images taken from the right in the top view panels as indicated by red arrowheads. Note that thalli are twisted in plants treated with TDIF, resulting in the uplift of the thalli from the medium as indicated by arrows.(B) Overall morphology of 14-day-old plants grown from gemmae. MpCLE1*ox* indicates overexpression under Mp*YUC2* promoter (proMpYUC2:MpCLE1) in Tak-1 and Mp*cik-1*^ge^ background. Mp*CLE1(1-417)ox*, indicates a truncate version of Mp*CLE1*. **(C)** Ground cover in 14-day-old plants (mean and SD; n= 11-14). Means sharing the superscripts are not significantly different from each other in Tukey’s HSD test, p < 0.05. Scale bars represent 1 cm in (A) and (B).

## Discussion

In this study, we performed functional analysis of Mp*CIK*, the sole *M. polymorpha* ortholog of Arabidopsis *CIK* genes, and showed that MpCIK is essential for MpCLE2 peptide signaling to regulate the apical meristem activity in gametophyte. Biochemical analysis in *N. benthamiana* supports the idea that the MpCLV1-MpCIK is a receptor-coreceptor pair for MpCLE2 peptide perception. Since the same ligand-receptor-coreceptor relationship of CLV3-CLV1-CIK is consistently observed for multiple paralogs in Arabidopsis (Hu et al., 2018; Cui et al., 2018; Anne et al., 2018), this system is likely an evolutionarily conserved mechanism for the perceotpion of CLV3-type CLE peptide in land plants. In addition, we suggested partial involvement of MpCIK in signaling of TDIF-type CLE peptide, MpCLE1, although MpCIK is not necessary for MpCLE1 signaling. Studies in Arabiodpsis have suggested that subclass-II receptors in the SERK subgroup function as coreceptors for TDIF-PXY/TDR signaling (Zhang et al., 2016b). Further studies on MpSERK would clarify the contribution of different coreceptors. The phylogenetic analysis inferred the divergence of CIK and SERK subgroups in the common ancestor of land plants, which coincides with the appearance of subclass-XI genes. Recent studies have also shown that SERK-interacting receptors from subclass X, such as BRI1/BRL and EMS1, are also encoded in bryophytes (Ferreira-Guerra et al., 2020; Furumizu and Sawa 2021). Interestingly, the two sequences from *Spirogyra pratensis* (Figure 1B) showed high similarity to the subclass-II genes from land plants. Studies on Zygnematales algae would provide a clue to understand the evolution of these receptors.

Gemma cups are specialized structures for vegetative propagation, found in certain species of Marchantiopsida (Yasui et al., 2019; Kato et al., 2020). Gemma cup formation initiates at the cells in the dorsal epidermis close to the apical meristem (Suzuki et al., 2020). We show that gain-of-function of Mp*CLE2* results in the delay of gemma cup formation although it is still unclear if the phenotypes in gemma cup formation can be uncoupled from the defects in the apical meristem. More detailed experiments will clarify the role for CLE in gemma cup formation. It is known that hormonal and environmental cues affect the formation of gemma cup (Flores-Sandoval et al., 2015; Aki et al., 2019; Li et al., 2020; Rico-Reséndiz et al., 2020). A possible role for MpCLE2 peptide signaling would be to mediate certain environmental cues to control the timing of gemma cup formation and thus clonal propagation, cooperatively with other hormonal inputs. In addition, involvement of CIK subgroup members into antiviral responses has been suggested in Arabidopsis (Fontes et al., 2004). Further studies of Mp*cik* knockout plants under various environmental conditions would provide a new insight into signals that allowed plants to survive on land.

Our biochemical data reveals that MpCIK is capable of associating with MpCLV1 or MpTDR in an ectopic and transient expression system of *N. benthamiana*. Futhermore, weak associations observed for MpCIK-MpCLV1 as well as MpCIK-MpTDR indicates possible requirement of other components in MpCIK-containg complex formation. For instance, in a ligand-induced dimerizatiom model, receptor-coreceptor interaction can be induced by the perception of ligand at their ectodomains, which in turn allows for their kinase domains to transphosporylate and activate signaling (Jaillais et al., 2011; Hohmann et al., 2018; Perraki et al., 2018). Thus, ligand and/or other membrane receptors could be required for strong MpCIK-MpCLV1/MpTDR association. With its genetic simplicity in CLE signaling, *M. polymorpha* will be a nice experimental system to address this point in future studies.

## Materials and Methods

### Phylogenetic analysis

Gene sequences of land plants were retrieved from Phytozome v12.1 database (https://phytozome.jgi.doe.gov/pz/portal.html) except for those of *Arabidopsis thaliana* (https://www.arabidopsis.org/), *Picea abies* (http://congenie.org/) and *Marchantia polymorpha* (https://marchantia.info/). Sequences of charophycean algae were reported in Bowman et al. 2017, obtained from transcriptome databases for *Spirogyra pratensis* (http://www.ncbi.nlm.nih.gov/Traces/wgs/wgsviewer.cgi?val=GBSM01&search=GBSM01000000&display=scaffolds) and *Coleochaete orbicularis* (http://www.ncbi.nlm.nih.gov/Traces/wgs/wgsviewer.cgi?val=GBSL01&search=GBSL01000000&display=scaffolds). Gene IDs and the protein sequences are listed in Table S1. Predicted protein sequences were aligned in Clustal W (https://www.genome.jp/tools-bin/clustalw). We excluded ambiguously aligned sequence to produce an alignment of 297 amino acid characters in the conserved cytosolic domain. Bayesian analysis was performed using MrBayes 3.2.7 (Ronquist et al., 2012). Two runs with four chains of Markov chain Monte Carlo (MCMC) iterations were performed for 1,500,000 generations, keeping one tree every 100 generations. The first 25% of the generations were discarded as burn-in and the remaining trees were used to calculate a 50% majority-rule tree. The standard deviation for the two MCMC iteration runs was below 0.01, suggesting that it was sufficient for the convergens of the two runs. Convergence was assessed by visual inspection of the plot of the log likelihood scores of the two runs calculated by MrBayes (Gelman and Rubin, 1992). Character matrix used for the Bayesian phylogenetic analysis is provided in Data Sheet S1.

### Plant materials and growth conditions

*Marchantia polymorpha* male Takaragaike-1 (Tak-1) accession was used as wild type in this study. *M. polymorpha* plants were grown on half-strength Gamborg B5 medium (pH 5.5) solidified with 1.4% agar at 22 °C under continuous white light. *Nicotiana benthamiana* seeds were grown on BM2 soil (Berger) in a growth room at 23 °C under continuous LED light.

### Peptide treatment

Synthetic peptides used in this study were analytically pure and dissolved in 0.1% TFA (trifluoroacetic acid) solution as stock solutions. For MpCLE2 peptide treatment, approximately 20 mature gemmae were floated on 2 mL liquid M51C medium containing 2% sucrose supplemented with 3 µM MpCLE2 peptide (KEVHypNGHypNPLHN) or mock (TFA) solution, in 12-well plates as described previously (Hirakawa et al., 2020). For the TDIF treatment, gemmae were plated on half-strength B5 agar plates supplemented with 3 µM TDIF (HEVHypSGHypNPISN) or mock (TFA) solution as described previously (Hirakawa et al., 2019).

### Constructs

Primers and plasmids are listed in Tables S2 and S3. All plant transformation vectors were generated using the Gateway cloning system (Thermo Fisher Scientific, MA, USA). Gatway destination vectors are described in Kinoshita et al., 2010, Ishizaki et al., 2015 and Sugano et al., 2018, except for pMpGWB301-YUC2p, which was generated in this study. A 3032 bp DNA flagment of Mp*YUC2* promoter sequence franking the translation initiation site was PCR amplified from pENTR-proMpYUC2 vector (Hirakawa et al. 2020) with a primer pair of MpYUC2prom3k_F_InFusion_XbaI and MpYUC2prom_R_InFusion_XbaI, and cloned into the *Xba*I digestion site of pMpGWB301 using In-Fusion HD Cloning Kit (Takara Bio, Shiga, Japan). For construction of *pro*Mp*YUC2*:Mp*CLE1*, entry clones, pENTR-MpCLE1 and pENTR-MpCLE(1-417) (Hirakawa et al., 2019), were transferred to the pMpGWB301-YUC2p vector using Gateway LR Clonase II Enzyme mix (Thermo Fisher Scientific). For genome editing of Mp*CIK*, a guide RNA was designed at the first exon/intron junction of Mp7g14210 using CRISPRdirect (https://crispr.dbcls.jp/) (Naito et al., 2015) and the plasmid for genome editing was constructed according to Sugano et al. 2018. For the expression of epitope-tagged receptors in *N. benthamiana*, coding sequences of Mp*CIK*, Mp*CLV1* and Mp*TDR* were PCR amplified from *M. polymopha* cDNA and cloned into pENTR/D-TOPO vector. Resultant entry clones (pENTR-MpCIK, pENTR-MpCLV1 and pENTR-MpTDR) were transferred to pXCSG-3FLAG or pXCSG-3HS vector using Gateway LR Clonase II Enzyme mix (Thermo Fisher Scientific).

### Production of transgenic *M. polymorpha*

Transgenic *M. polymorpha* plants are listed in Table S3. *Agrobacterium*-mediated transformation of *M. polymorpha* was performed using regenerating thalli according to Kubota et al. 2013. CRISPR/Cas9-based genome editing was performed according to Sugano et al. 2018 and mutations in the guide RNA target loci were examined by direct sequencing of PCR product amplified from genome DNA samples with primers listed in Table S2. Genome editing of Mp*CLV1* was performed as described previously (Hirakawa et al., 2020). Nomenclature of genes and mutants are according to Bowman et al. 2016.

### Plant imaging and phenotypic measurement

Overall morphology of plants was obserbed under a digital microscope (DMS1000, Leica Microsystems, Wetzlar, Germany) or under a digital camera (TG-6, Olympus, Tokyo, Japan). For the quantification of ground cover area in plant images, blue color was extracted and quantified using ImageJ (Schneider et al., 2012). For the measurement of apical notch width, plants grown on liquid medium were individually transferred onto agar medium and imaged under a digital microscope (DMS 1000, Leica Microsystems). To quantify the apical notch width, distance between the rims of apical notch was measured on the obtained images using ImageJ (Schneider et al., 2012).

### Promoter GUS assay

Individual plants were stained separately in 30-50 µL GUS staining solution (50 mM sodium phosphate buffer pH 7.2, 1 mM potassium-ferrocyanide, 1 mM potassium-ferricyanide, 10 mM EDTA, 0.01% Triton X-100 and 1 mM 5-bromo-4-chloro-3-indolyl-β-D-glucuronic acid) at 37 °C in dark. GUS-stained samples were washed with water, cleared with ethanol and mounted with clearing solution for imaging under a light microscope (BX51, Olympus).

### Transient expression in *N. benthamiana*

Agrobacterium tumefaciens strains GV3101 MP90RK carrying expression constructs were grown in YEB medium with appropriate antiboiotics, harvested by centrifugation at 4,500 rpm for 10 min, and resuspended in infiltration buffer [10 mM MES (pH 5.7), 10 mM MgCl2, 150 μM acetosyringone]. The cultures were adjusted to an OD 600 of 1.0 and incubated at room temperature for at least 3 h prior to infiltration. Equal volumes of cultures of different constructs were mixed for co-infiltration, and then mixed with agrobacterial cultures (OD 600 of 1.0) carrying the p19 silencing suppressor in a 1:1 ratio (Voinnet et al., 1999). The resulting cultures were infiltrated into leaves of 3-to 4-week-old *N. benthamiana*. The leaf samples were harvested 3 d after infiltration for subsequent protein extraction (Betsuyaku et al., 2011).

### Protein extraction

Total protein was extracted from the infiltrated *N*.*benthamiana* leaves with IP extraction buffer (1: 1 w/v, 50 mM Tris–HCl pH 8.0, 150 mM NaCl, 10 % glycerol, 1 % Triton X-100, 1×Proteinase inhibitor cocktail SIGMA P9599 and 1 mM EDTA) and incubate the extract at 4 °C for 30 min. The lysates were centrifuged at 20,000×g for 20 min at 4 °C and the supernatants were then centrifuged again at 20,000×g for 5 min at 4 °C. The resultant supernatants were used as total protein lysates.

### Co-immunoprecipitation

For immunoprecipitaion, 1 ml of the lysates prepared with IP extracation buffer from 0.5 g of leaves was incubated with anti-HA Affinity Matrix (Roche 11815016) for o/n in a rotary shaker at 4 °C. The beads were collected and washed three times with 1 ml of the extraction buffer. Immunopricipitaed proteins were eluted from the beads by boiling in SDS sample buffer at 95 °C and analyzed by Western blot using the corresponding antibodies. We used the following antibodies; Anti-HA-Peroxidase High Affinity (3F10) (Roche 12013819001) and Monoclonal ANTI-FLAG M2-Peroxidase (HRP) (SIGMA A8592).

## Data Availability Statement

The original contributions presented in this study are included in the article/Supplementary Material.

## Author Contributions

YH conceived the study, designed the work with input from all authors, and prepared the manuscript draft. GT, SB, NO and YH performed the experiments and analyzed the data. All authors contributed to the article and approved the submitted version.

## Acknowledgments

We thank colleagues in the peptide/protein center of WPI-ITbM for peptide synthesis, Eriko Betsuyaku and Ikuko Nakanomyo for technical assistance.

